# Silencing of a raspberry homologue of *VRN1* is associated with disruption of dormancy induction and misregulation of subsets of dormancy-associated genes

**DOI:** 10.1101/2024.02.13.580140

**Authors:** Brezo Mateos, Katharine Preedy, Linda Milne, Jenny Morris, Pete E. Hedley, Craig Simpson, Robert D. Hancock, Julie Graham

## Abstract

Winter dormancy is a key process in the phenology of temperate perennials. The changing climate is severely impacting its course leading to economic losses in agriculture. A better understanding of the underlying mechanisms, as well as the genetic basis of the different responses, are necessary for the development of climate-resilient cultivars. This study aims to provide an insight into winter dormancy in red raspberry (*Rubus idaeus* L).

We report the transcriptomic profiles during dormancy in two raspberry cultivars with contrasting responses. The cultivar ‘Glen Ample’ showed a typical perennial phenology, whereas ‘Glen Dee’ registered consistent dormancy dysregulation, exhibiting active growth and flowering out of season. RNA-seq combined with weighted gene co-expression network analysis (WGCNA) highlighted gene clusters in both genotypes that exhibited time-dependent expression profiles, Functional analysis of ‘Glen Ample’ gene clusters highlighted the significance of the cell and structural development prior to dormancy entry as well the role of genetic and epigenetic processes such as RNAi and DNA methylation in regulating gene expression. On the contrary, dormancy release in ‘Glen Ample’ was associated with upregulation of transcripts associated with the resumption of metabolism, nucleic acid biogenesis and processing signal response pathways.

Many of the processes occurring in ‘Glen Ample’ were dysregulated in ‘Glen Dee’ and twenty-eight transcripts exhibiting time-dependent expression in ‘Ample’ that also had an Arabidopsis homologue were not found in all samples from ‘Glen Dee’. These included a gene with homology to Arabidopsis *VRN1* (*RiVRN1.1*) that exhibited a sharp decline in expression following dormancy induction in Glen Ample. Characterisation of the gene region in the ‘Glen Dee’ genome revealed two large insertions upstream of the ATG start codon. We propose that non-expression of a specific VRN1 homologue in ‘Glen Dee’ causes dormancy misregulation as a result of inappropriate expression of a subset of genes that are directly or indirectly regulated by *RiVRN1.1*.

**HIGHLIGHT:** The raspberry cultivar Glen Dee exhibits aberrant winter dormancy status associated with insertions in the upstream promoter region of a *VRN1* (*RiVRN1.1*) homologue that silence expression, allowing the identification of dormancy-associated genetic modules that are regulated by *RiVRN1.1*.

## INTRODUCTION

Winter dormancy is an adaptive mechanism protecting temperate plants from abiotic stress. It is established at the end of summer as a response to shortening photoperiods and falling temperatures. Once the onset of dormancy is established, the plants require exposure to a genotype-dependent time at cold temperatures to allow resumption of growth. Dormancy induction is triggered by environmental cues, particularly daylength and temperature, which in many species exhibit an interaction that impacts not only the onset of growth cessation and bud formation but also the depth of bud dormancy (Olsen 2010). Climate change is therefore decoupling the main environmental cues for dormancy induction and the rise in average winter temperatures is compromising the fulfilment of cold requirements for dormancy release. This is leading to alterations in the phenology of dormancy, uneven budbreak, and frost damage. Subsequently, natural ecosystems as well as several temperate crops are being negatively impacted (Amano et al. 2010; Cleland et al. 2007; Fitter 2002; Ford et al. 2016; Mosedale et al. 2016; Tixier et al. 2019).

Red raspberry *Rubus idaeus* (L.) is a perennial berry crop belonging to the family Rosaceae. It is primarily cultivated across temperate areas, restricted mainly by the need for winter chill. *R. idaeus* is a diploid species with a relatively small genome (~300 Mb), although highly heterozygous (Price et al. 2023). Many of its commercial varieties have a biennial life cycle, requiring dormancy to fully transition to a reproductive stage. However, there are annual genotypes able to flower within the first year of growth. Cold exposure is still needed for optimum and consistent flowering (Foster et al. 2019). Both the type of cycle and the extent of the cold requirements have been key breeding targets in the species (Jennings 1988). Frost damage and inconsistent budbreak have become increasingly frequent, causing significant economic losses (Graham and Simpson 2018; Heide and Sønsteby 2011; Sønsteby and Heide 2014).

Raspberry dormancy research has been limited by the difficulty in phenotyping dormancy depth and the genetic resources available. Several QTL studies in biparental populations from annual x biennial cultivars have tackled the inheritance of the type of cycle. A single locus conferring annual fruiting was described in tetraploid blackberry (*Rubus* subgenus *Rubus* Watson.) (Castro et al. 2013), although its transferability to raspberry genetic maps has not been possible (Foster et al. 2019). Two QTL *(RiAF3* and *RiAF4*) associated with annual fruiting have been mapped in chromosomes 3 and 4 respectively (Jibran et al. 2019) of red raspberry. Candidate genes underlying QTL were proposed based on their function and differential expression. A homolog of *JMJ14 (JUMONJI14)*, encoding a H3K4 demethylase was identified in the region of *RiAF3*. *PFT1 (PHYTOCHROME AND FLOWERING TIME 1), FCA (FLOWERING CONTROL LOCUS A),* and *AGL24 (AGAMOUS-LIKE 24)* were the candidates proposed for *RiAF4*. More recently, several loci linked to annual cycle were described on chromosomes 1, 2, 4, 5, and 6 (Graham et al. 2022).

Mazzitelli et al. (2007) explored the molecular mechanisms underlying dormancy release in raspberry in an early RNA microarray analysis. This study reported a significant abundance of genes involved in stress mechanisms throughout the entire process. An *SVP*-like MADS-box gene *RiMADS_01* was identified, showing a profile of constant downregulation as dormancy was released. *RiMADS_01* was subsequently mapped to a region on chromosome 5 that showed significant association with flowering time in a biparental population exhibiting differences in phenology (Graham et al. 2009). Interestingly, *RiMADS_01* is homologous to *dam6* of *Prunus persica*, proposed as a negative regulator of bud break (Jiménez et al. 2010).

A growing body of research has unveiled some of the main mechanisms involved in dormancy induction and release in other species. RNA-seq analysis, sometimes combined with epigenetic or metabolomic analysis have contributed significantly to this area. However, some fundamental questions regarding signalling, underlying causes of the differences in cold requirements between genotypes, and the conservation of general mechanisms across taxa remain unsolved. Here, we report a comparative analysis of dormancy induction and release aimed to provide an insight into the processes occurring during dormancy in two raspberry genotypes with contrasting cold requirements, Glen Ample and Glen Dee. We identify transcriptional misregulation in Glen Dee associated with poor aberrant dormancy regulation. Genetic analysis revealed the presence of insertions in the promoter of *VRN1*-like gene in Glen Dee that silence its expression, thereby identifying genetic modules associated with bud dormancy in raspberry that are dependent on *VRN1*-like expression.

## MATERIALS AND METHODS

### Plant material and dormancy assessments

The material for this study was collected from mature plants of Glen Ample, a high chilling requirement cultivar (Mazzitelli et al. 2007) and Glen Dee, which we have previously identified as a low chilling requirement cultivar. The plants were kept under commercial conditions in a polytunnel at The James Hutton Institute, in Invergowrie (56°27’24.9”N 3°03’56.1”W), Scotland. Plants were pruned, leaving 2 to 3 canes per root (stool), individually tagged. Tissue sampling was conducted fortnightly between 6th August 2021 and 14th March 2022. At each time point, four canes of each cultivar were randomly selected, covering the length of the tunnel. The axillary buds from the middle and top region of each cane were pooled and flash frozen in liquid nitrogen. All the material was collected between 10 am and 12 noon to standardise the circadian variation.

The progression of dormancy was monitored through single-node tests (Velappan et al. 2022). Five canes of each cultivar were cut into nodes and transferred to a forcing environment. The nodes were suspended in water trays at 20 °C with light regimes of 16 h above 100 Wm^−2^ for 14 days after which the number of dormant, dead, and active buds was recorded.

### RNA sequencing and development of the reference transcriptome

Total RNA was isolated from the bud tissue using Qiagen RNeasy kit according to the manufacturer’s instructions. Quality was checked using a Bioanalyzer 2100 (Agilent) prior to generating individual indexed libraries each from 1 μg RNA using the Stranded mRNA Prep kit (Illumina), as recommended. Libraries were quality controlled and pooled in equimolar quantities. The final pool was run at 750 pM with 2% PhiX control library on a NextSeq 2000 (Illumina) with a P3 300 cycle kit, according to manufacturer guidelines, to generate 150 bp paired-end data. Data was demultiplexed, generating individual fastq files for each sample. A reference transcriptome was developed using the methodology described in Coulter et al. (2022). RNA-seq reads were then mapped onto a raspberry reference transcript dataset (RTD) using SALMON (Patro et al. 2017) to produce transcripts per million (TPM) quantifications.

### Network construction

Clusters of genes with similar expression patterns were identified using Weighted CO-expression Network Analysis (WGCNA). Raw data from SALMON quantification was transformed into length-scaled TPM at gene level using the R package tximport (Soneson et al. 2015). Genes within the 5^th^ percentile of variance were filtered, lowering the input from 37297 to 34941. Reads were normalised using the variance stabilising transformation (vst) from the package DESeq2 (Love et al. 2014). Samples were clustered based on Euclidean distance to detect potential outliers. The soft threshold for the network construction was estimated based on the fit of the linear model regressing *log[p(k)] log(k)*, where p(k) corresponds to the frequency distribution of the connectivity of the nodes. Six was used as consensus power for both genotypes. Co-expression networks were built for each genotype using the R package WGCNA (Langfelder and Horvath 2008). Modules whose overall expression changed over time were identified by fitting a model using the function lmFit from the package limma (Ritchie et al. 2015), the eigengene of each cluster as response, and time point as predictor. Standard errors were smoothed using empirical Bayes (eBayes) from limma. Results were filtered using 0.001 as significance threshold.

The main processes represented in every cluster were identified through functional profiling using g:GOSt in g:Profiler (Kolberg et al. 2023). *Arabidopsis thaliana* was used as reference with the analysis run using default settings (s:SCS threshold algorithm for multiple testing correction, user threshold of 0.05).

### Comparative analysis

Comparative analysis focused on the four clusters of genes upregulated during dormancy induction in Glen Ample (M1, M31, M55, and M9). Reads of both genotypes for each cluster were clustered applying k-means method with two centres. Genes without reads in Glen Dee were identified; from them *RiVRN1.1* was analysed further.

### Sequencing *RiVRN1.1*

The sequence of *RiVRN1.1* was blast searched against the reference genome of Glen Moy (Hackett et al. 2018), giving a unique hit in scaffold 3118. The genomic sequence of the gene plus a fragment of around a 1 Kb at both ends was selected. Sequences of genomic DNA and mRNA were aligned using the MAFF algorithm in Benchling. Introns, exons, and 3’ and 5’ UTRs were identified. Primers within the 5’ and 3’ were designed using NCBI’s primer design tool. Their specificity was verified through BLAST against the Glen Moy genome.

Sequencing primers were designed from the exons in the mRNA of Glen Ample and used to sequence the coding region of both genotypes using Sanger sequencing. For sequencing the promoter and UTR regions, an extra set of primers was designed using the reference genome of Glen Moy.

### Sequencing the genomes of Glen Ample and Glen Dee

High-quality genomic DNA (250 ng each) for cultivars Glen Ample, Glen Mor and Glen Dee was used to generate indexed sequencing libraries using the Illumina DNA Prep kit, as recommended. Following library QC on a Bioanalyzer 2100 (Agilent), libraries were pooled, loaded at 750 pM with 1% PhiX control, and sequenced as paired-end 150 bp reads on a NextSeq 2000 (Illumina) using a P1 sequencing kit, according to the manufacturer’s instructions. Fastq files were demultiplexed post-run on the NextSeq prior to analysis. Each genomic DNA data were mapped onto the Anitra genome using BWA-MEM (Li 2013). The SAM output was run through ‘sambamba view’ (Tarasov et al. 2015) using a filter to remove mappings with more than 6 mismatches to the reference. The output BAM files were sorted and had duplicate reads removed using samtools (Danecek et al. 2021) and sambamba respectively.

Contigs containing RiVRN1 in the genomes of Glen Moy, Glen Mor, Glen Ample, and Glen Dee were aligned using the MAFF algorithm in Benchling. The promoter region of Glen Ample and Glen Dee was confirmed through PCR and Sanger sequencing of the amplicon.

## RESULTS

### Winter phenology of Glen Ample and Glen Dee

This study monitored the progression of dormancy in two genotypes on a bi-weekly basis between August 2021 and March 2022. The depth of dormancy was measured as the proportion of bud break after 14 days in a forcing environment. The genotypes analysed had contrasting chilling requirements, previously estimated as 1500 chilling hours for Glen Ample (Mazzitelli et al. 2008), and 750 for Glen Dee (Sutherland et al. 2019). Glen Ample exhibited a rapid change in bud dormancy status, shifting from almost entirely active buds on 5^th^ August to its maximum dormancy on 30^th^ September (TP5), with ~65% of excised buds remaining dormant (**Figure 1a**). Endodormancy was almost entirely released by 10^th^ November (TP8). Glen Dee exhibited a shallower entry into dormancy, with the majority of buds remaining active until 16^th^ September. After this date, an increasing proportion of buds exhibited a dormant state, reaching a maximum dormancy on 15^th^ October (TP6). However, even at maximum dormancy only about 35% of buds were dormant. As was observed for Glen Ample, dormancy was released in the first half of November (TP8), although in some canes a higher proportion of dormant buds were observed in late November and early December (TP9 and 10), before almost all buds resumed activity for the remaining samples. A key observation was that isolated buds from Glen Dee showed a much larger dormancy distribution than those isolated from Glen Ample, as can be seen from the distribution of replicates from the four canes. The dispersion of the measurements suggests that dormancy is a relatively synchronous process within individuals of Glen Ample. In contrast, canes of Glen Dee varied up to between 12 and 81% of dormant buds at TP7. This seems particularly acute around the onset of dormancy. Indications that developmental processes were more widely perturbed were supported by the observation that flowering also appears to be dysregulated in Glen Dee with flowers observed on field planted canes as late as 13^th^ December (**Fig. 1b**). On the contrary, Glen Ample canes appeared entirely dormant in the field with closed buds (**Fig. 1c**).

**Figure 1.**
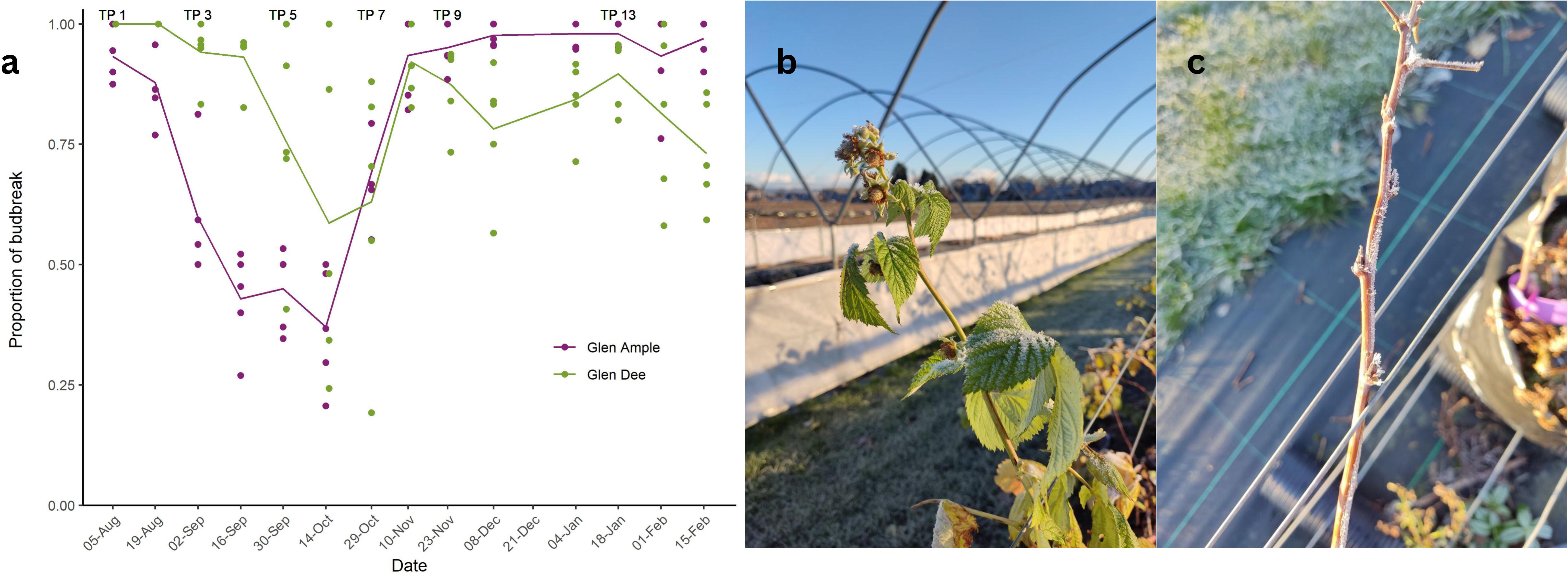
**a-**Profiles of dormancy depth in axillary buds of raspberry cultivars Glen Dee and Glen Ample, sampled between August 2021 and February 2022. Dormancy depth was measured as the proportion of budbreak in node sections exposed to a forcing environment for 14 days. Each point corresponds to a cane (replicate). **b-**Canes of Glen Dee exhibiting active growth and flowering on 12 December 2022, suggesting a dysregulation of dormancy consistent over seasons. **c-**Canes of Glen Ample on 12 December 2022. This genotype showed a *wild type* dormancy response: the plants are senescent; the buds are covered with scaly leaves and no flowering was registered.

### Co-expression networks

To understand the mechanisms underpinning differences in dormancy behaviour between the two genotypes, RNA-seq was undertaken. RNA was isolated from bud tissue of a subset of six time points representing different dormancy status (**Fig. 1a**) and sequenced using Illumina. Transcript abundance was quantified using SALMON. The raspberry RTD contains 137902 transcripts that mapped to 37492 unique genes. The length-scaled transcripts per million (TPM) for every gene was estimated for the 48 samples. Genes whose variance fell within the 5th general percentile were filtered, leaving 34941. Given the highly contrasting dormancy behaviours between the genotypes (**Fig. 1a**) the dataset was split for the analysis, and an independent network was built for each genotype. The network of Glen Ample clustered the genes into 64 modules containing between 110 and 7962 genes that differed in their expression profiles. Each module’s profile was summarised in an eigengene and 2472 genes were assigned to the grey module (not assigned to any specific expression profile). The network built from Glen Dee samples comprised 56 clusters ranging in size from 117 to 6082 genes and 1966 genes were assigned to the grey module. Interestingly, both networks include two clusters with opposite profiles and a notably higher size: modules 1 and 2 of Glen Ample (7962, 5694 genes), and modules 1 and 2 of Glen Dee (6082, 6068 genes). To identify clusters that exhibited changes in transcript abundance over time, a limma model where ModuleEigengene I] time was fitted. Nine modules in the network of Glen Ample and 5 of Glen Dee were significant (p<0.001) (**Fig. 2**, **3**). Hub genes, highly connected nodes within each of the significant clusters, were identified. Supplementary table S1 lists the top five hub genes for each module.

**Figure 2.**
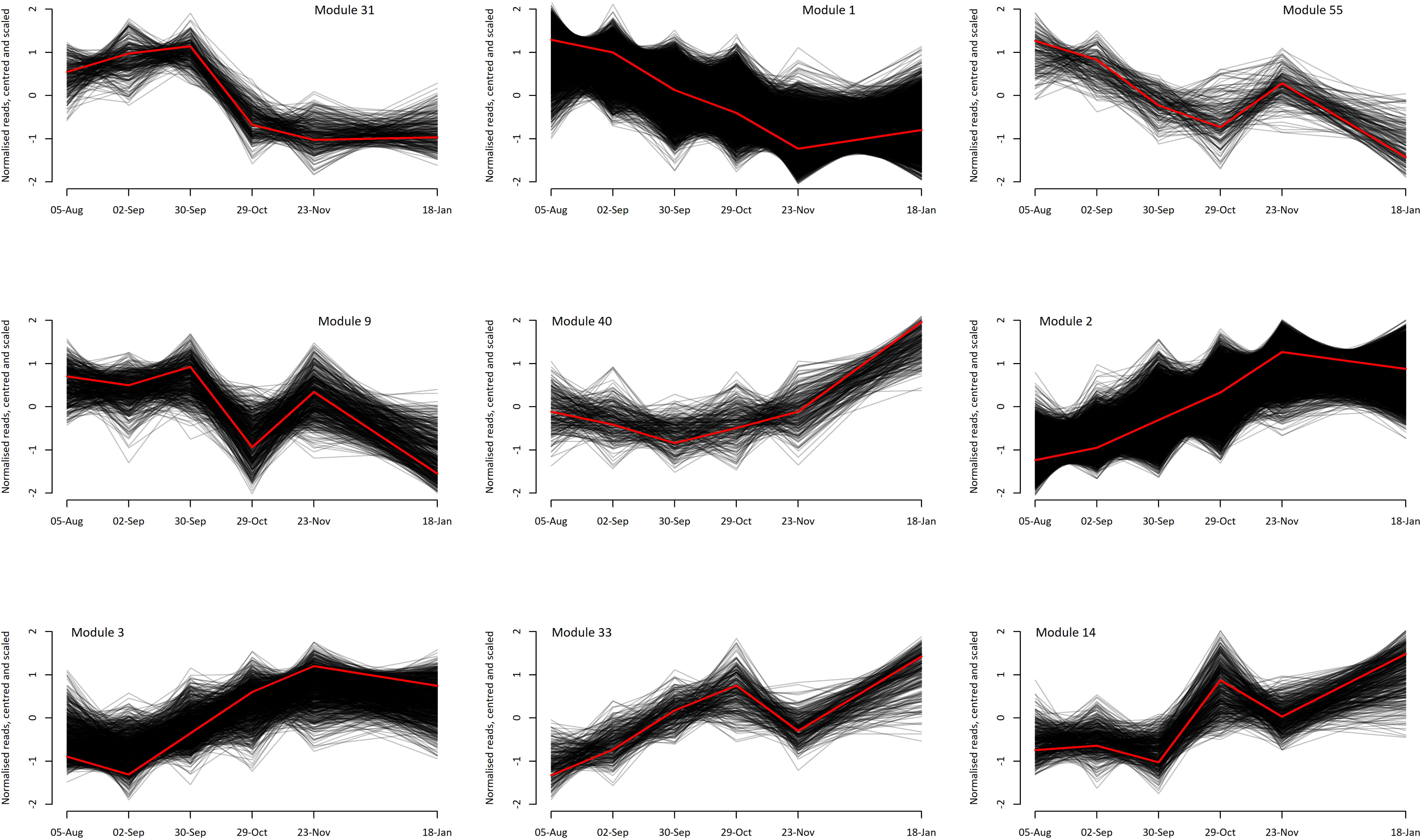
Gene expression in clusters differentially expressed in Glen Ample. WGCNA methodology produced a network of clusters of co-expressed genes. Clusters of interest fitted a limma model using time as predictor (p< 0.001). Each cluster is identified by an eigengene (red) summarising the profile of the genes within. Gene expression values are represented as TPM centred and scaled.

**Figure 3.**
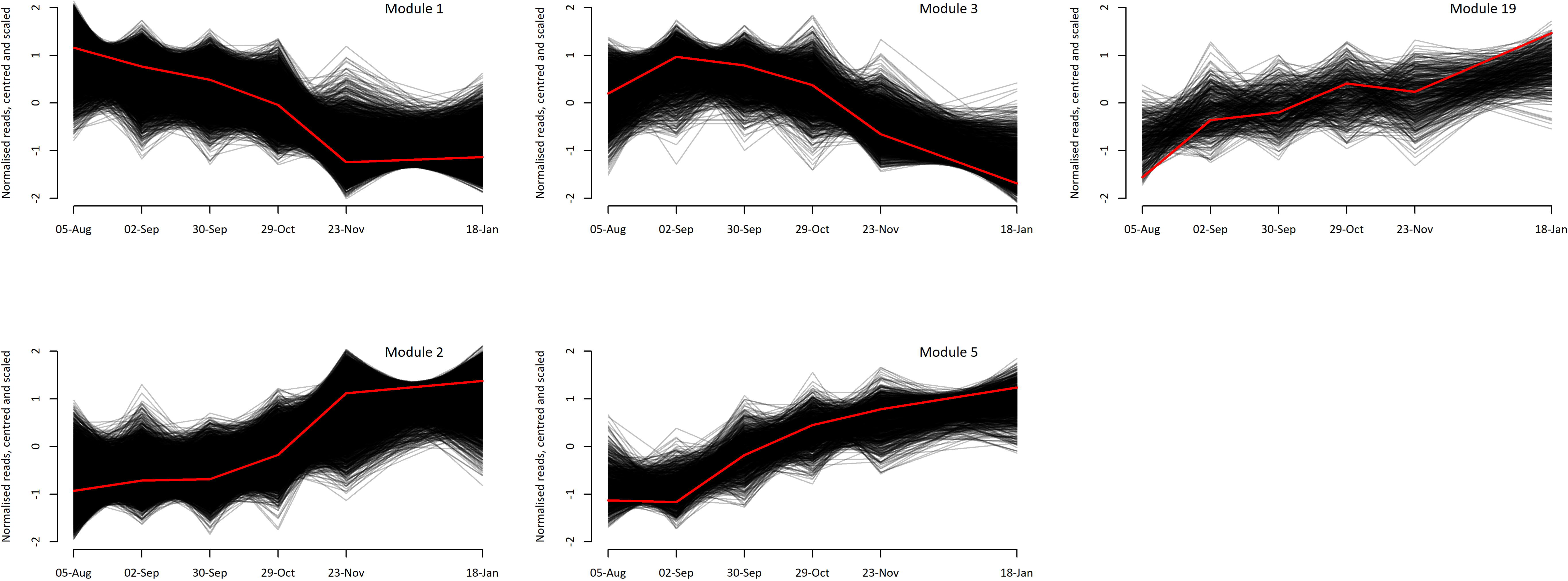
Gene expression in clusters differentially expressed in Glen Dee. WGCNA methodology produced a network of clusters of co-expressed genes. Clusters of interest fitted a limma model using time as predictor (p< 0.001). Each cluster is identified by an eigengene (red) summarising the profile of the genes within. Gene expression values are represented as TPM centred and scaled.

The network of Glen Ample is composed of 9 significant differentially expressed clusters. They can be grouped into two main branches, clusters that contain transcripts that show a pattern of decline over time (M1, M31, M55 and M9) and those that contain transcripts exhibiting an increase in abundance over time (M40, M2, M3, M33 and M14). The first branch consists of four clusters (M1, M31, M55, M9) comprising transcripts abundant at the end of summer and progressively decreasing as dormancy is established (**Fig. 2**). This branch can be linked to mechanisms active within dormancy induction. Clusters M1 and M55 had the peak of expression in the first time point, corresponding to 5^th^ August, and decreased in abundance falling to a low level around the onset of dormancy (TP 5, 30/9/2021). M1 showed a constant reduction, reaching its minimum at the point of release (TP9, 23/11/2021). Contrastingly, the abundance of M55 rose slightly at that point. These two clusters were defined as Induction I subphase of dormancy as their maximum expression was at the earliest time point and consistently declined as dormancy status increased. Clusters M9 and M31 show high relative abundances during induction until TP5, when plants reach a high depth of dormancy. Both exhibited a sharp reduction toward the release (TP7 29/10/2021). Phenologically, these mechanisms correspond to a second subphase of dormancy induction (Induction II).

Clusters M33 and M14 registered low abundances throughout dormancy induction and peak during dormancy release (TP7 29/10/2021). Both modules decreased at the end of dormancy release (TP9, 23/11/2021) to reach maximum abundance in active buds (TP13 18/1/2021). These two modules have been grouped as a subphase (release I). M2 shows low levels of relative expression at induction, increasing consistently to a maximum at the end of dormancy release (TP9, 23/11/2021) and decreasing towards TP13 (18/1/2021). Genes within M3 exhibit low abundances during early induction, increasing gradually towards release. Interestingly, the pattern of expression of this cluster is highly inconsistent between replicates. Finally, M40 clusters genes with low abundancy throughout dormancy induction and release, peaking sharply at TP 13 (18/1/2021) once buds are fully responsive.

The network of Glen Dee can be similarly divided in two main branches of clusters, showing continuous upregulation and downregulation respectively (**Fig. 3**). The first branch groups M1 and M3 with high abundance during induction and decreasing towards release. Genes in M1 have maximum abundance at TP1 (5/8/2021), while M3 peaks at TP 3(2/9/2021). The transition between the two main phases occur at TP 7 (29/10/2021), a month later than Glen Ample. The clusters M19, M2, and M5 follow a similar profile, registering low abundance in the first time points and increasing expression over time. M5 and 19 reach maximum expression at TP 13 (18/1/2021) after endodormancy is released. M2 experiences a sharp increase at TP9 (23/11/2021) that continues at TP13 (18/1/2021). The structure of the network contrasts with results from Glen Ample, which showed a greater level of resolution and consistency between replicates, implying a loss of some degree of control.

### Functional analysis of gene clusters

Functional profiling using g:GOST allowed the identification of the main biological processes represented by every cluster. This, combined with the month-to-month resolution of the networks, provided a timeline for dormancy induction and release. As Glen Ample established and released dormancy successfully and consistently, this network was used as reference.

#### Dormancy induction I

This subphase includes genes in clusters M1 and M55, with maximum expression at the earliest time point and downregulated towards dormancy onset. Cluster M1 groups 7962 genes, 6177 of them have an associated TAIR homologue. g:GOST analysis revealed 14 driver terms regarding biological processes (Supplementary table 2). Highest enrichment was detected for the GO term cell cycle processes (p = 3.74E-39), including functions such as cell division (mitosis), development of anatomical structures and microtubule-based processes, suggesting active growth. In addition, organization or biogenesis of cell wall, morphogenesis of plant epidermis, microtubule-based movement (p = 3.38E-12), and formation of xylem and phloem (p = 0.001), and protein transport were enriched. GO terms for metabolism were highlighted as drivers, including metabolism of lipids (p = 3.63E-17), aromatic amino acids, lactone (p = 0.018), and phenylpropanoids (p = 2.49E-12). The main regulatory processes significantly enriched in this cluster were DNA methylation (p = 0.020) and histone lysine methylation (p = 0.005), as well as regulation of hormone levels (p = 0.047). Genes involved in RNAi-mediated antiviral immune response were significantly overrepresented (p = 0.003).

Cluster M55 is considerably smaller, containing 166 genes of which 136 are annotated. The genes in M55 express similarly to M1, having maximum expression in August at early stages of dormancy induction and decreasing at dormancy onset. However, this cluster shows a small increase in expression towards the release (TP 9), although inconsistent between replicates. All the enriched GO terms in this cluster are related to membrane transport processes and endoplasmic reticulum localization.

#### Dormancy induction II

The second subphase corresponds to genes that reach maximum expression at later stages of induction. Module M31 groups 267 genes, of which 223 have available annotation. g:GOST analysis showed significant enrichment in genes involved in carbohydrate catabolism (p = 0.007), cell wall modification (p = 0.016), and biosynthesis and metabolism of phenylpropanoids (p = 0.046, p= 0.007). This cluster groups genes exhibiting high abundances during dormancy induction whose expression increase until maximum at the point of dormancy onset, falling in expression afterwards. Cluster M9 grouped 421 genes annotated from a total of 509. Functional analysis identified four significantly enriched terms, vesicle-mediated transport (p = 1.88E-10), protein catabolic process (p = 5.898E-06), protein K63-linked ubiquitination (p = 0.045), and vacuolar proton-transporting V-type ATPase complex assembly (p = 0.049). This suggests strong protein turnover during this stage, a likely consequence of intense gene expression during development of protective structures at earlier stages of dormancy.

#### Dormancy release I

The earliest processes occurring during dormancy release correspond to clusters M33 and M14, and M3, with 260, 434, and 947 genes, respectively, from which 204 and 268, and 604 are annotated. No significantly enriched GO terms were detected for cluster M14. Cluster M33 showed overrepresentation for cellular process (p = 0.012), transport (p = 0.026), and sulphur compounds biosynthetic processes (p = 0.0332). Although not highlighted as a driver term, establishment of location within the cell was also significantly overrepresented (p = 0.037). Cluster M3 is upregulated during all processes of release, but inconsistent between replicates. From significantly overrepresented GO terms, three were highlighted as drivers: secondary metabolism (p = 7.566E-05), protein phosphorylation (p= 8.65E-07), and response to stress (p = 3.16E-17).

#### Dormancy release II

A second subphase of dormancy release peaks at TP9, at the last stages of release. This contains cluster M2, a big cluster containing 5694 genes, 4675 of which have annotation available, and M3, smaller (947 genes, 604 annotated) and quite variable in profile among the samples. Genes in M2 registered low relative abundance during dormancy induction, increasing continuously to a maximum at TP9, and decreasing towards active tissue (TP13). g:GOST analysis provided 248 significantly enriched terms regarding biological processes detailed in Supplementary table 2, 14 of which were highlighted as driver terms. The majority of driver terms are related to resumption of gene expression, such as mRNA metabolic process (p = 1.81E-21), nitrogen compound transport (p = 1.79E-10) and response to organonitrogen compound (p = 1.054E-6), organophosphate biosynthesis, DNA-templated transcription initiation (p = 0.008), cytoplasmic translation (p = 0.009), translational initiation (p = 0.010), and protein folding (p = 3.154E-05). Aligned to this, post-embryonic development (p = 6.74E-23) appeared as a driver term. Starch catabolism (p = 0.0367) also appears as a driver term, suggesting a role as energy source.

#### Resumption of growth

Cluster M40 exhibited a peak in gene expression at the latest sampled timepoint once dormancy was fully released. This cluster is composed of 231 transcripts of which 171 genes are annotated. Functional profiling highlighted non-coding RNA (ncRNA) processing (p = 4.045E-05), macromolecule modification (p = 0.008), and metabolic processes, as well as nitrogen compound and organic cyclic compounds metabolism specifically (p = 0.005, p = 0.019), all presumably associated with resumption of growth.

### Comparative analysis of Glen Dee and Glen Ample

The overall mechanisms dysregulated during dormancy induction in Glen Dee were investigated using the gene network of Glen Ample as reference. As Glen Dee is unable to fully establish dormancy, analysis focused on the four clusters of genes upregulated during dormancy induction in Glen Ample. For each cluster, reads of both genotypes were merged and reclustered using K-means with two centres. This split the reads into a centre containing reads from Glen Ample and genes of Glen Dee with similar profiles, and a second centre containing genes with altered profiles in Glen Dee. gOST analysis from gProfiler identified ontology terms overrepresented as potential dysregulated mechanisms (Supplementary table 3). During early induction cell cycle, microtubule-based process and movement, lipid, lactone, and phenylpropanoids, and histone lysine methylation showed altered expression. Contrastingly with Glen Dee, these mechanisms did not shift to downregulation towards the maximum of dormancy but stayed upregulated for much of the experiment. In addition to these mechanisms, other highlighted processes such as photosynthesis (p=0.03683) and stomatal movement (p=0.0405) were unique to this genotype. Dysregulated genes in module 55 were significantly enriched in genes associated with the endosome (GO:0005768, p = 0.034379) and those in module 31 were significantly enriched in genes associated with apoplast, chloroplasts and plastids (GO:0048046, GO:0009536, GO:0009507). Terms for synthesis of phenylpropanoids and flavonoids were also overrepresented (WP:WP1538, GO:0009698; p = 0.011596, p = 0.011054), as well as copper ion binding, NADP+ binding, and oxidoreductase activity (GO:0005507, GO:0070401, GO:0016491; p = 0.017675, p = 0.024927, p = 0.025497). Re-clustering module 9 did not produce a separation, but rather split reads of Glen Ample into two clusters approximately even, and reads of Glen Dee followed a similar pattern.

Genes not expressed in Glen Dee were identified as potential candidates regulating dormancy in Glen Ample. Module 55 of Glen Ample contains 166 genes of which 3 were not detected in Glen Dee. These genes were not annotated in the reference transcriptome. Module 9 of Glen Ample contains 506 genes, 10 of which were not detected in Glen Dee. Four of them have alignments in BLAST, but their Arabidopsis homologues are proteins of unknown function (AT5G45530.1 and AT2G36430.1), and the other two have a hAT dimerisation domain (AT5G33406.1), and an FBD-like domain (AT1G51055.1) but no known function. Module 31 of Glen Ample contains 267 genes, 4 of which were not detected in the transcriptome of Glen Dee. None produced significant alignments. Module 1 of Glen Ample contains 7962 genes, of which 148 not detected in Glen Dee samples. Twenty-four of these transcripts had homologues in Arabidopsis as determined by significant BLAST hits (Table 1).

**Table 1.** Genes in cluster M1 of Glen Ample not detected in Glen Dee. Genes were annotated using Arabidopsis as reference. M1 groups genes with maximum levels of expression at early dormancy induction, decreasing expression throughout the remainder of the time course.

| Gene | Homolog | Description |
| --- | --- | --- |
| rirt3_HiC_scaffold_1100G000080 | AT2G01050.1 | Sinc ion binding, nucleic acid binding |
| rirt3_HiC_scaffold_1118G000160 | AT3G17080.1 | Plant self-incompatibility protein S1 family |
| rirt3_HiC_scaffold_1G002370 | AT3G42170.1 | BED zinc finger, hAT family dimerisation domain |
| rirt3_HiC_scaffold_1G020110 | AT5G45520.1 | Leucine-rich repeat (LRR) family protein |
| rirt3_HiC_scaffold_1G021000 | AT4G08685.1 | SAH7 Pollen Ole e 1 allergen and extensin family protein |
| rirt3_HiC_scaffold_1G026130 | AT3G18990.1 | VRN1 REM39 AP2/B3-like transcriptional factor family protein |
| rirt3_HiC_scaffold_270G000220 | AT4G03460.1 | Ankyrin repeat family protein |
| rirt3_HiC_scaffold_2G010200 | AT2G21340.1 | MATE efflux family protein |
| rirt3_HiC_scaffold_2G014650 | AT5G53130.1 | CNGC1 ATCNGC1 cyclic nucleotide gated channel 1 |
| rirt3_HiC_scaffold_2G047170 | AT1G64185.1 | Lactoylglutathione lyase / glyoxalase I family protein |
| rirt3_HiC_scaffold_330G000110 | AT1G65780.1 | P-loop containing nucleoside triphosphate hydrolases superfamily protein |
| rirt3_HiC_scaffold_3G038200 | AT1G08390.1 | unknown protein RecQ-mediated genome instability protein 2 |
| rirt3_HiC_scaffold_3G039040 | AT4G14060.1 | Polyketide cyclase/dehydrase and lipid transport superfamily protein |
| rirt3_HiC_scaffold_4G010850 | AT5G57590.1 | BIO1 adenosylmethionine-8-amino-7-oxononanoate transaminases |
| rirt3_HiC_scaffold_4G010900 | AT1G17230.1 | Leucine-rich receptor-like protein kinase family protein |
| rirt3_HiC_scaffold_5G007100 | AT2G34730.1 | myosin heavy chain-related |
| rirt3_HiC_scaffold_5G016800 | AT4G12500.1 | Bifunctional inhibitor/lipid-transfer protein/seed storage 2S albumin superfamily protein |
| rirt3_HiC_scaffold_5G040920 | AT5G36930.2 | Disease resistance protein (TIR-NBS-LRR class) family |
| rirt3_HiC_scaffold_5G045300 | AT1G27170.1 | transmembrane receptors ATP binding |
| rirt3_HiC_scaffold_6G038650 | AT4G32208.1 | heat shock protein 70 (Hsp 70) family protein |
| rirt3_HiC_scaffold_775G000010 | AT1G54690.1 | HTA3 H2AXB G-H2AX GAMMA-H2AX gamma histone variant H2AX |
| rirt3_HiC_scaffold_7G001240 | AT5G24280.1 | GMI1 gamma-irradiation and mitomycin c induced 1 |
| rirt3_HiC_scaffold_7G002170 | AT5G24280.1 | GMI1 gamma-irradiation and mitomycin c induced 1 |
| rirt3_HiC_scaffold_7G009310 | AT3G53960.1 | Major facilitator superfamily protein Peptide/nitrate transporter plant |

### Characterising *RiVRN1.1*

From the 28 genes upregulated during dormancy in Glen Ample and not expressed in Glen Dee, rirtd3_HiC_scaffold_1G026130 was analysed further. This gene is annotated as *VRN1*, *VERNALIZATION1* in *Arabidopsis thaliana*. *VRN1* is involved in vernalisation and transition between vegetative and reproductive stages, and is widely characterised in Arabidopsis (reviewed in Banerjee, Wani, and Roychoudhury 2017), as well as barley (Deng et al. 2015) and wheat (reviewed in Milec, Strejčková, and Šafář 2023). This gene homologue was highly expressed in Glen Ample during dormancy induction up to time point 3 (**Fig. 4**), but transcripts sharply declined at TP5 and remained close to the limits of detection for the remainder of the time course. There are 4 splice variants, with rirtd3_HiC_scaffold_1G026130.RTD.1 being the longest and most abundant transcript. Alignment of the mRNA sequences of Glen Ample with the reference genome of Glen Moy (Hackett et al. 2018) showed no mismatches.

**Figure 4.**
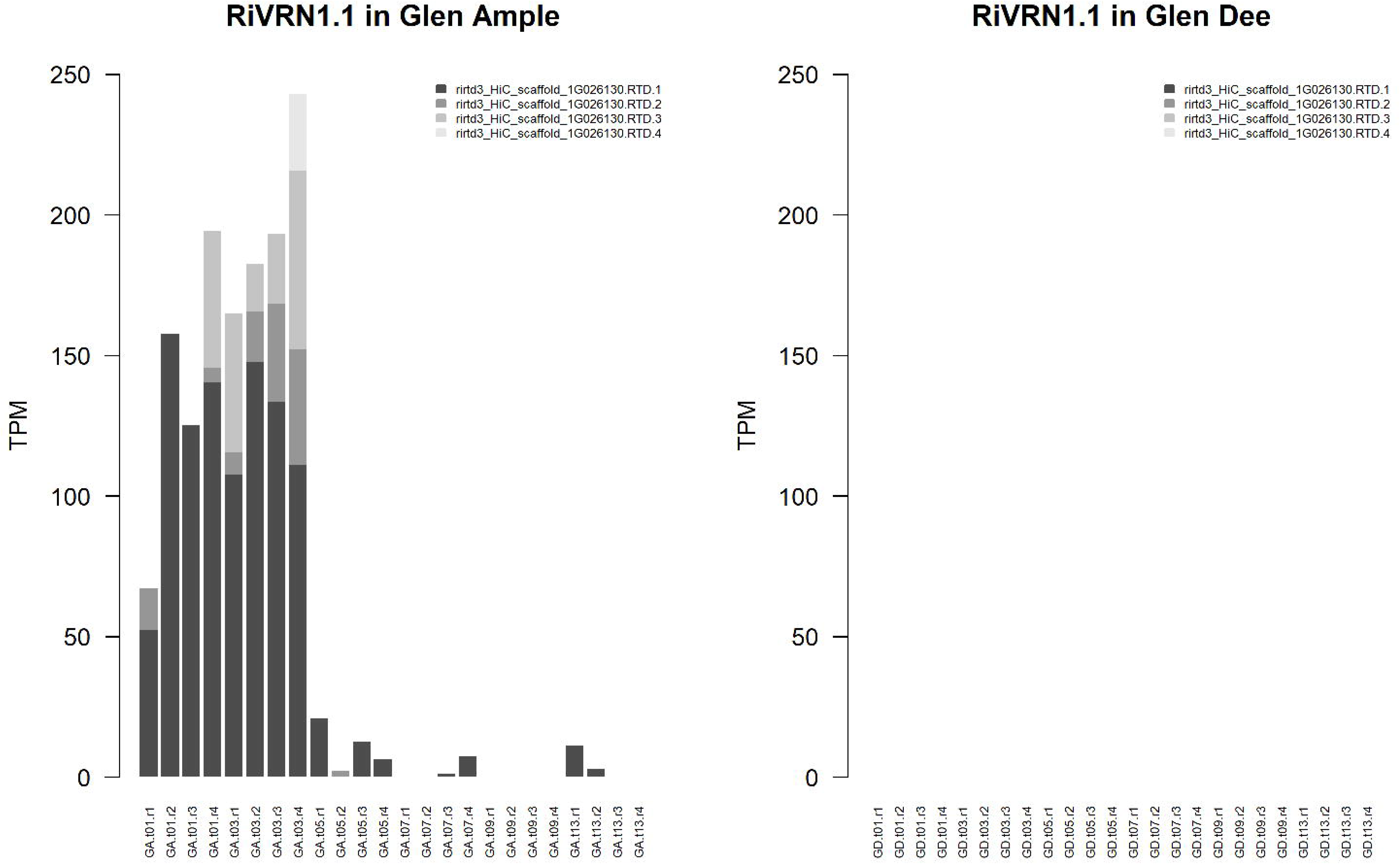
Expression of *RiVRN1.1* (rirtd3_HiC_scaffold_1G026130) in Glen Ample and Glen Dee samples. Four isoforms were detected in Glen Ample, reaching the maximum accumulated expression at TP 3 (2 September 2021). No reads of any *RiVRN1.1* transcript were detected in samples from Glen Ample.

The *RiVRN1.1* isoforms of Glen Ample were aligned to identify the splicing sites. The coding region of Glen Ample and Glen Dee was re-sequenced using Sanger sequencing. Although substitutions were detected, most mutations are synonymous with Glen Moy, and no new stop codons or shifts in reading frame were observed in the Ample and Dee genomes relative to Glen Moy. The genomes of Glen Ample and Glen Dee were sequenced using Illumina to access the promoter regions and gene environment. Alignment of contigs containing rirtd3_HiC_scaffold_1G026130 revealed two insertions in the promoter region of the gene in Glen Dee. The most distal insertion is 747 bp in length and is located 535 bp upstream of the mRNA (**Supplementary material 1**). Sequence and presence of the insertion were confirmed through amplicon sequencing. The second insertion is 301 bp and located 54 bp upstream of the beginning of the mRNA in Glen Dee (**Fig. 5**). These insertions likely disrupt the promoter, leading to silencing of rirtd3_HiC_scaffold_1G026130.

**Figure 5.**
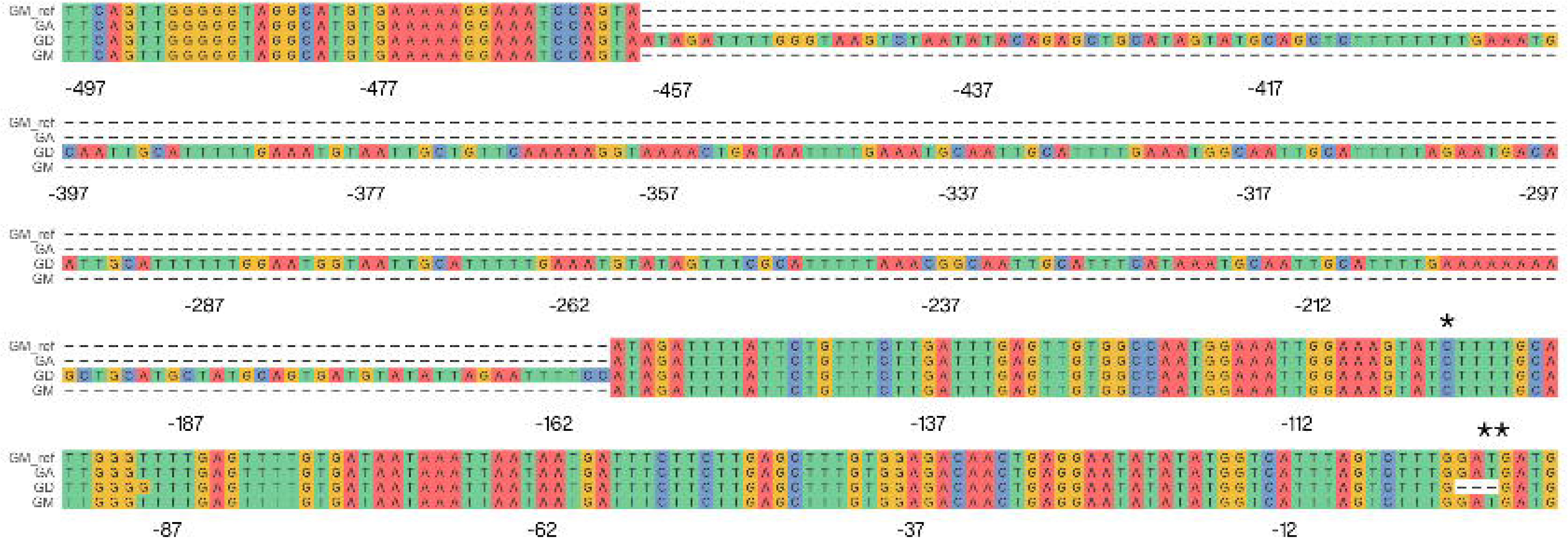
Alignment of genomic sequences of Glen Moy (reference genome and PCR product), Glen Ample, and Glen Dee, illustrating the second insertion located in the promoter area of *RiVRN1.1*. The PCR product from *RiVRN1* of Glen Ample, Glen Moy, and Glen Dee was Sanger sequenced and aligned with the reference genome of Glen Ample. Glen Moy and Glen Ample showed a high level of conservation. Glen Dee has a 301 bp insertion 54 bp upstream of the start of the UTR of Glen Ample (*) and 162 bp upstream of the start codon (**). In addition, Glen Dee has a substitution 93 bp upstream the start codon, and a deletion of three bases where Glen Ample and Glen Moy have two consecutive ORFs.

Twelve more genes show homology with *VRN1* of *Arabidopsis thaliana* (**Supplementary material 2**) and their expression levels vary during dormancy, suggesting their cues and functions may differ. Alignment of their mRNA sequences showed relatively low identities, ranging from 61.95 to 86.5%, to the predominant isoform of rirtd3_HiC_scaffold_1G026130 (rirtd3_HiC_scaffold_1G026130.RTD.1).

## Discussion

We investigated the genetic mechanisms underlying dormancy in two raspberry cultivars with contrasting phenology. Reportedly, Glen Ample and Glen Dee differ in their chilling requirements, which would translate in different timings of dormancy release. Bi-weekly monitoring of the dormancy status revealed differences not only in the timing or the cold requirements, but in the profile of dormancy and consistency among different individuals. In addition, blooming out of season was recorded in Glen Dee as late as 12^th^ December 2021.

The onset and release of dormancy occurred earlier than expected based on the literature in both genotypes. This may be due to most of the published data being assessed in whole canes, therefore incorporating effects of paradormancy. In our experiment, endodormancy was released before the end of winter and subsequent frost risk.

### A model for dormancy in *Rubus idaeus*

The time-resolved co-expression network of Glen Ample was used to build a model for dormancy in raspberry. The process is divided into two main phases, induction and release, controlled by independent mechanisms. Induction initiates at the end of summer, in early August, and completes late September.

Induction can be divided into two subphases. Its earliest stage, Induction I, involves metabolic, cellular organisation and gene silencing processes. Key metabolic processes include the organisation of the cell wall or biogenesis, as well as potentially linked processes such as metabolism of phenylpropanoids, or aromatic amino acid metabolic processes, and their precursors. Primary metabolic processes are represented by lipid metabolism. During this stage histone lysine methylation, DNA methylation, and RNAi-mediated immune response genes are overrepresented. These processes likely initiate silencing of gene expression that would lead to growth arrest. Involvement of histone and DNA methylation in the establishment of winter dormancy has been widely described in other temperate species (Rothkegel et al. 2020; W. Chen et al. 2022; Conde et al. 2013; Ríos et al. 2014). The role of RNAi mechanisms in the process is less studied, although variations in the miRNA linked to QTL controlling chilling requirements have been reported in peach (Barakat et al. 2012). Interestingly, homologues of some of the genes detected in our analysis, such as *AGO1* (*ARGONAUT 1)* or *HEN1* (*HUA ENHANCER 1*) play a central role in the silencing of gene expression during seed dormancy in *Arabidopsis thaliana* (Jones-Rhoades and Bartel 2004; Allen et al. 2005; Tognacca and Botto 2021).

A second group of mechanisms peak a month later, around the time the plants reach the midpoint of dormancy (Induction II). Cell wall modification and linked processes, such as phenylpropanoid metabolism are active. General metabolism is represented by catabolism of carbohydrates. Other processes include vesicle-based transport, membrane docking, organelle localisation by membrane tethering and processes associated with endoplasmic reticulum (ER) organisation (GO:0051643, GO:2261817, GO:0090158). The potential significance of the latter may relate to the ER being an integral component of the plasmodesmata (Nicolas et al. 2017) which have been strongly implicated in dormancy induction and release (Rinne et al. 2011).

At this stage, processes linked to protein turnover are strongly represented in the cells. This could be a consequence of growth and development of protective structures occurring in earlier stages and could potentially have an additional regulatory role, as has been proposed during the release of seed dormancy (Oracz and Stawska 2016). This group of genes have a second peak in expression about the time of dormancy release.

Onset of dormancy coincides with a shift in mechanisms in the transcription data. Release of dormancy was a gradual process, less synchronous between individuals than induction. The earliest stage, named here Release I, contains genes upregulated at the maximum levels of dormancy. The main processes include secondary metabolism, phosphorylation of proteins, and abiotic stress signalling, particularly hypoxia. Mazzitelli et al. (2007) reported several genes involved in stress response linked to dormancy in raspberry. Resemblance of gene expression in dormant buds and response to anoxic stress has been previously drawn (Considine and Foyer 2014) and proposed to be involved in release through increase in ROS (Beauvieux et al. 2018). Here, this group of genes shows low relative abundances through dormancy onset and induction, peaking sharply during release. Interestingly, timing of upregulation and maximum abundance seems to be inconsistent between replicates (**Fig. 2**) but does not reflect the consistency in the release of dormancy at corresponding time points (**Fig 1a**). Interestingly, this GO term includes *CBP1 (CCG-BINDING PROTEIN 1)*, *EXL2 (EXORDIUM-LIKE 2), ATRMA3, EDF1 (ETHYLENE RESPONSE DNA BINDING FACTOR 1),* and *CYP707A3 (CYTOCHROME P450),* reported in gene regulatory networks linked to dormancy (Tarancón et al. 2017).

After the initial stages of anoxic stress, endodormancy is released between October and late November. Gene expression is resumed as is inferred by overrepresentation of genes involved in transcription, translation, and protein folding. Starch catabolism is one of the main processes occurring at this stage, potentially an energy source for cell machinery resumption after the period of low carbon input from short photoperiods. However, a growing body of evidence suggest a pivotal role of non-structural carbohydrates in dormancy release signalling (Tixier et al. 2019, 2018; Gibon et al. 2004; Bolouri Moghaddam and Van den Ende 2013; Palacio et al. 2014). This has been hypothesized to rely on the effect of temperature on the enzymatic equilibrium between starch and sugars (Tixier et al. 2019). Our data shows a constant increase in expression of this group of genes from the onset of dormancy to the resumption of growth, following the accumulation of chilling. Further analysis is needed to clarify the role of starch degradation as cause or consequence of release of dormancy. Upon resumption of growth, a group of genes involved in regulation of non-coding DNA and different aspects of metabolism become upregulated, overlapping with the processes previously described.

### *RiVRN1.1* is a key to the establishment of dormancy

In our data, disruption of the promoter region of *RiVRN1.1* is associated with reduced dormancy and early flowering, confirmed over two following seasons. *RiVRN1.1* registers high relative abundances during dormancy induction in Glen Ample. After reaching maximum mid-induction, expression falls abruptly, remaining at low abundance for the rest of the time course. The cultivar Glen Dee exhibits two insertions of 774 bp and 300 bp, at the distal and proximal regions of the promoter, and no expression was detected in our data. *RiVRN1.1* is an ortholog of *VRN1,* (*VERNALIZATION 1),* a MADS-box transcription factor holding a central role in the vernalization pathway in *Arabidopsis thaliana* by repressing *FLC* (*FLOWERING LOCUS C*) (Levy et al. 2002; Milec et al. 2023). Links between regulation of vernalization and dormancy through epigenetic mechanisms have been previously reviewed (Considine and Foyer 2014; D. Horvath 2009; Maurya and Bhalerao 2017; Ríos et al. 2014).

In addition to the flowering and dormancy phenotypes, the gene expression data of Glen Dee shows mechanisms involved in photosynthesis, active growth, and stomatal movement, active throughout the winter. Therefore, dormancy induction seems to be lacking a key point of regulation. Comparative analysis, taking the network of Glen Ample as a model, identified processes altered during dormancy induction in Glen Dee. Most processes identified in the ontology analysis are clustered in the same module as *RiVRN1.1.* This includes regulatory elements, such as hormones, histone lysine methylation, DNA methylation, and RNAi silencing.

The activity of *VRN1* in Arabidopsis has been linked to changes in the methylation patterns of histone H3 (Bastow et al. 2004). Several genes altered in Glen Dee from the histone lysine methylation pathway are involved in silencing mechanisms, such as *CHROMOMETHYLASE 3 (CMT3), ARGONAUTE 4 (AGO4)*, *PDP1*, *ATRFC, KRYPTONITE (KYP)*, and *SDG34*. *AGO4* was proposed to act as repressor of dormancy in wheat seeds (Singh et al. 2013; Katsuya-gaviria et al. 2020). In transcriptomic studies in poplar, leafy spurge, and tea tree, *AGO4* shows downregulation during bud endodormancy (D. P. Horvath et al. 2008; Matzke and Mosher 2014; Howe et al. 2015; Hao et al. 2017). *KYP* encodes a methyltransferase known to hold a role in seed dormancy by regulating negatively *DOG1* and *ABI3* (Zheng et al. 2012). Genes regulating flowering time were present in the cluster of altered expression profiles. These include *EARLY FLOWERING MYB PROTEIN (EFM)*, *RBBP5 LIKE (RB), EIN6 (ETHYLENE INSENSITIVE 6)* and *TRAUCO (TRO)*, as well as regulators of *FLC*, *EARLY FLOWERING IN SHORT DAYS (EFS)* (Kim et al. 2005), *PLANT HOMOLOGOUS TO PARAFIBROMIN* (Park et al. 2010), *PDP1 (PWWP DOMAIN PROTEIN 1)* (Zhou et al. 2018), and *VERNALIZATION INDEPENDENCE 3 (VIP3)* (Zhang et al. 2003). *RBBP5* is a component of the *COMPASS* complex, and its silencing supresses the activity of *FLC*, leading to early flowering (Jiang et al. 2011). *EIN6 (REF6)* inhibits dormancy through the catabolism of ABA in seeds of Arabidopsis (H. Chen et al. 2020). *EFM* is a key repressor of *FT* integrating light and temperature stimuli (Yan et al. 2014). Interestingly, *EFM* expression is promoted by *SVP (SHORT VEGETATIVE PHASE),* homologous to the *DAM (DORMANCY-ASSOCIATED MADS-BOX)* family.

In addition to the constitutive silencing of *RiVRN1.1*, other mechanisms and genes could be contributing to the dysregulation of dormancy observed in Glen Dee. Chromatin accessibility through histone methylation is one of the central processes controlling dormancy induction (W. Chen et al. 2022). An ortholog of *H2A.X* was found among the genes involved in dormancy induction in Glen Ample and not transcribed in Glen Dee (Table 1). This gene encodes a gamma-induced variant of histone H2A in *Arabidopsis thaliana*. *h2a.x* mutants have been linked to tissue-specific hypomethylation in the endosperm of Arabidopsis (Frost et al. 2023). However, its mechanism of action remains unsolved and, to our knowledge, has not been studied in vegetative meristems.

## Conclusions

This study provides an overview of the mechanisms underlying raspberry dormancy. The WGCNA methodology identifies changes in gene expression on a month-by-month basis. Comparative analysis of transcriptomic and phenological data from Glen Ample and Glen Dee identified a general dysregulation of dormancy induction in the latter, which leads to growth and flowering out of season. Analysis of transcriptomic and genomic data identified a *VRN1-*like gene (*RiVRN1.1*) as the likely candidate for these responses. The genome of Glen Dee exhibits two insertions in the proximal and distal promoter regions of *RiVRN1.1* that might cause disruption and silencing. Our findings provide a framework for molecular analysis of dormancy in raspberry and suggest the role of *RiVRN1.1* as key point in the regulation of dormancy induction.

## Acknowledgements

Brezo Mateos acknowledges the Mylnefield trust for receipt of a PhD scholarship. Work was funded by the Mylnefield Trust and the Rural and Environmental Science and Analytical Services Division of the Scottish Government under the 2022-2027 Strategic Research Programme.

## Table legends

**Supplementary table 1.** Top hub genes of the differentially expressed gene clusters of Glen Ample and Glen Dee. Hub genes are genes highly connected within each cluster. The table summarises the top 5 for each differentially expressed cluster and genotype.

**Supplementary table 2.** Summary of the functional analysis of the gene clusters differentially expressed in Glen Ample. Source refers to the type of term (GO: Gene Ontology, BP: Biological Process); term_id represents the GO code. Highlighted identifies the terms highlighted as drivers (Kolberg et al. 2023), the p values of the overrepresentation test are provided adjusted and as negative log10 of the adjusted p-value. Size parameters correspond to the described by Kolberg et al. (2023) for the overrepresentation test. The query size corresponds to the number of annotated genes within each cluster submitted for the functional analysis. Intersections identifies the genes on each query belonging to a given GO term.

**Supplementary table 3.** Summary of the functional analysis of the of dysregulated genes in Glen Dee corresponding to clusters 1, 31, and 55 of Glen Ample. Source refers to the type of term (GO: Gene Ontology, BP: Biological Process); term_id represents the GO code. Highlighted identifies the terms highlighted as drivers (Kolberg et al. 2023), the p values of the overrepresentation test are provided adjusted and as negative log10 of the adjusted p-value. Size parameters correspond to the described by Kolberg et al. (2023) for the overrepresentation test. The query size corresponds to the number of annotated genes within each cluster submitted for the functional analysis. Intersections identifies the genes on each query belonging to a given GO term.

